# Differential Effects of Posttranslational Modifications on the Membrane Interaction of Huntingtin Protein

**DOI:** 10.1101/2023.12.01.569526

**Authors:** Zhidian Zhang, Charlotte Gehin, Luciano A Abriata, Matteo Dal Peraro, Hilal Lashuel

## Abstract

Huntington’s disease (HD) is a neurodegenerative disorder caused by an expanded polyglutamine stretch near the N-terminus of the huntingtin (HTT) protein, rendering the protein more prone to aggregate. The first 17 residues in HTT (Nt17) interact with lipid membranes and harbor multiple posttranslational modifications (PTMs) that can modulate HTT conformation and aggregation. In this study, we used a combination of biophysical studies and molecular simulations to investigate the impact of PTMs on the helicity of Nt17 in the presence of various lipid membranes. We demonstrate that anionic lipids such as PI4P, PI(4,5)P2, and GM1 significantly enhance the helical structure of unmodified Nt17. This effect is attenuated by single acetylation events at K6, K9, or K15, whereas tri-acetylation at these sites abolishes Nt17 membrane interaction. Similarly, single phosphorylation at S13 and S16 decreased but did not abolish the POPG and PIP2-induced helicity, while dual phosphorylation at these sites markedly diminished Nt17 helicity, regardless of lipid composition. The helicity of Nt17 with phosphorylation at T3 is insensitive to the membrane environment. Oxidation at M8 variably affects membrane-induced helicity, highlighting a lipid-dependent modulation of Nt17 structure. Altogether, our findings reveal differential effects of PTMs and crosstalks between PTMs on membrane interaction and conformation of HTT. Intriguingly, the effects of phosphorylation at T3 or single acetylation at K6, K9, and K15 on Nt17 conformation in the presence of certain membranes do not mirror that observed in the absence of membranes. Our studies provide novel insight into the complex relationship between Nt17 structure, PTMs and membrane binding.

**TOC figure:** 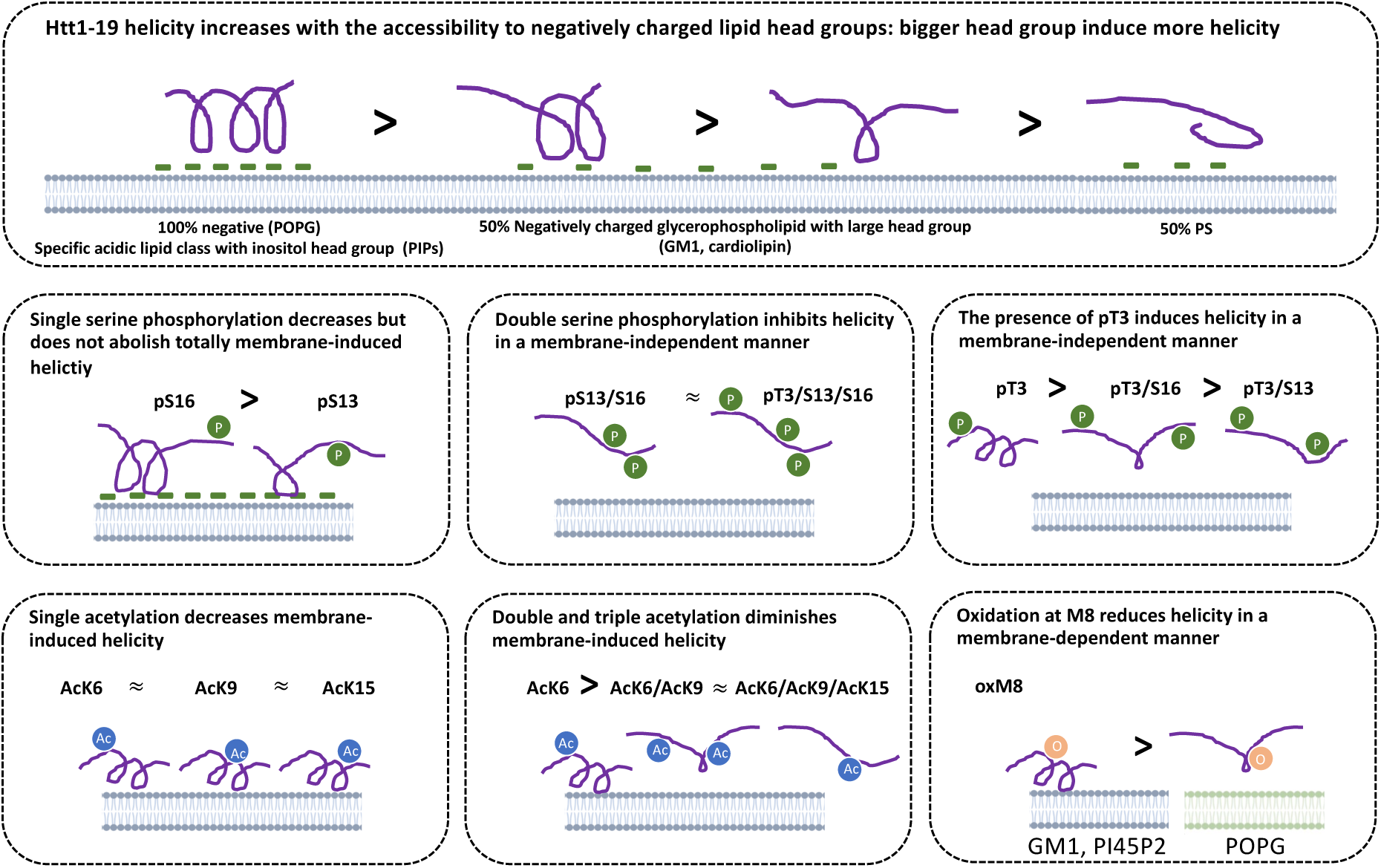

## Introduction

Huntington’s disease (HD) is an inherited fatal neurodegenerative disease caused by the aberrant expansion of the CAG triplet repeat sequence within the first exon of *huntingtin* gene (*IT15*) (Macdonald 1993). In HD patients, a mutant form of the Huntingtin (mHTT) harboring the expanded polyglutamine stretch (polyQ ≥ 36) renders the protein more prone to aggregate and form cytoplasmic and nuclear inclusions ^1–3^.

Converging evidence point to an N-terminal fragments containing exon1 as the primary species responsible for HTT aggregation and inclusion formation in HD ^4^. Among the various N-terminal huntingtin fragments found in the brain, overexpression of mutant forms HTT exon 1 (Httex1) with expanded polyQ repeats has been shown to be sufficient to recapitulate many features of HD pathology in cellular and animal models of HD ^5,6^. Unlike other N-terminal fragments, Httex1 is naturally produced via aberrant splicing in a CAG-repeat dependent manner ^6^. Altogether, these findings suggest that mHttex1 plays important roles in the pathogenesis of HD.

Httex1 is composed of a central polyglutamine (polyQ) domain, flanked by a N-terminal 17-residue amphipathic stretch (Nt17) and 50-residue proline-rich (PR) domain in C-term. In solution, Httex1 is intrinsically disordered but can adopt a tadpole-like conformation, and the compaction of this conformation, aggregation, and pathogenicity of Httex1 is influenced by several factors, including the polyQ repeat length ^7,8^, post-translational modifications (PTMs)^9–11^, pH ^12^, or interactions with membranes ^13^.

Several PTMs localized in the Nt17 domain have been shown to modulate the structure ^9,14–16^, aggregation ^10,11,15,17–21^, toxicity ^11,18,22,23^, clearance ^11,22,24^, nuclear translocation ^24,25^, and membrane interaction ^19,20,26,27^. A decreased level of phosphorylated T3 was found in neuronally differentiated induced pluripotent stem cell (iPSC) lines from HD patient ^17^. Phosphorylation of mutant Httex1 at S13 and S16 mediated by TBK1 (TANK-binding kinase 1) decreases Httex1 aggregation and inclusion formation in different cellular models and a C. elegans model of HD ^28^. The introduction of phosphometics at S13 and S16 reversed the pathology of mutant HTT in an HD mouse model ^18^. Furthermore, the M8 oxidation level is increased in the R6/2 HD mouse model ^29^. These experimental observations suggested that PTMs play important roles in regulating mutant Httex1 pathogenic properties.

Recent advances in semisynthetic synthesis ^9,14^ combined with in vitro kinase phosphorylation studies ^24^ enabled mechanistic studies to decipher the impact of each single PTM site as well as their crosstalk effect in Nt17 domain. Phosphorylation at S13 and/or S16 inhibits mutant Httex1 aggregation and decrease the helicity of Nt17 in vitro. Phosphorylation at T3 increases the helicity of Nt17 and inhibits aggregation of mutant Httex1, while acetylation of single lysine residues (K6, K9, or K15) does not impact Httex1 aggregation. The effects of phosphorylation on helicity and aggregation are reversed by the acetylation at K6 but not other acetylation sites in Nt17 ^9^. Also, while AcK6 alone has no effect on aggregation, crosstalk between AcK6 and oxidation at M8 drastically suppresses HTT aggregation ^15^. Altogether, these findings suggest that the crosstalks between PTMs within Nt17 play important roles in the regulation of Httex1 structure and aggregation.

Both the full-length HTT and its N-terminal fragments can associate with membranes and vesicles in the brain of healthy individuals and HD patients ^1,30,31^. For example, both they were found to interact with synaptosomes, endoplasmic reticulum, and mitochondrial membranes in cultured cells ^13,32^. Some studies have suggested that the aberrant interactions between mHTT and membranous organelles could disrupt ER and nuclear envelope ^33–35^. Httex1 anchors to membranes through its Nt17 domain that form an amphipathic helix at the surface of the lipid bilayer ^13,36,37^.

Our understanding of the sequence and structural determinants of Httex1 interactions with membranes remains incomplete. PTMs at Nt17 influence Httex1-membrane interaction as well as Httex1’s structure and aggregation. For example, chemical acetylation of all three lysine residues K6, K9, and K15 by sulfo-N-hydroxysuccinimide was shown to inhibit membrane interactions ^26^. Phosphomimetic mutations S13D/S16D inhibit membrane binding and membrane-mediated aggregation ^38^. Atomistic simulations showed that HTT Nt17 interacts with phospholipid in four main steps: approach, reorganization, anchoring, and insertion ^39^. *In vitro,* the binding properties of HTT and the conformation of the Nt17 domain are sensitive to the lipid composition ^31,37,40–42^, surface charge and curvature ^38,43^ of the membrane models. Even though mutant Httex1 can self-assemble in solution, several studies have also shown that its misfolding and aggregation could also be catalyzed by the presence of membranes. Mutant Httex1 with expanded polyQ repeat (e.g. 46Q) exhibit enhanced fibrillization in the presence of a membrane composed of anionic lipids POPG and POPS even at low lipid-to-protein ratios^44^. The profound influence of membrane lipids in HD was also highlighted by the impaired lipid metabolism of gangliosides in fibroblasts of HD patients and disrupted ganglioside levels in caudate samples from human HD subjects ^45–47^. Finally, altered sphingomyelin fatty acid composition was found in cerebral white matter in HD patients ^48,49^.

Research on Nt17 interactions with membranes has only explored limited lipid composition diversity and was mostly performed using PTM-mimicking mutation or non-site-specific chemical modifications and focused on investigating one PTM at a time ^26,38^. Thus, a systematic study of the HTT-membrane interaction with membranes of diverse lipid compositions as well as HTT Nt17 with different combinations of site-specific PTMs is essential for assessing the impact of lipid composition and PTM crosstalk on HTT-membrane interaction. Towards this goal, we used a combination of biophysical and atomistic molecular dynamics (MD) approaches to investigate Nt17 interactions with brain lipid extract enriched with different lipids and with physiological membrane mimetics. We also investigated how PTMs influence Nt17 interactions with membranes of different compositions using Httex1 proteins bearing *bona fide* N-terminal acetylation (specific acetylation at K6, K9 and K15), single and multiple phosphorylations at T3, S13 and S16, and oxidation at M8. These studies enabled us to gain new insights and atomistic-level understanding of the complex interplay between PTMs, Nt17 structure, and lipid composition in regulating Httex1 interaction with biological membranes.

## Results and Discussion

### The presence of membranes influenced the helicity of unmodified and PTM’ed Nt17/19

To assess the effect of PTMs on the membrane interaction of unmodified and PTM’ed Nt17 (Table S1), we prepared 17 types of large unilamellar vesicles (LUVs), either made of brain lipid extract (TBLE) and 10 or 50%mol of lipids implicated in the interaction of huntingtin with membranes (Table S2), or made of lipid compositions that mimicked the membrane of different organelles (Table S2-S4). Next, we compared the change of helicity of modified and unmodified Nt17 or Nt19 (Nt17 + QQ) in the presence of membranes. We used both Nt17 and Nt19. Previously, we showed that there is no difference in the overall structure between Nt17 and Nt19 ^9^. We used circular dichroism (CD) to measure changes in the conformation of Nt17 as an indirect indication of interaction between HTT and membrane, because previous studies showed with NMR experiments that Nt17 undergoes a conformation transition to a helical form upon membrane association ^37,38,50^. The effects of AcK6, AcK9, and AcK15 on Nt19 conformations were independent of the presence of membranes, except for GM1 and PI45P2 (Figure 1, Figure S1-S2). Similarly, the effect of pT3, pT3pS13, pT3pS13pS16, and pT3pS16 did not change in the presence of most membranes, except in the presence of GM1, PI45P2, PI4P, cardiolipin, and POPG.

**Figure 1.**
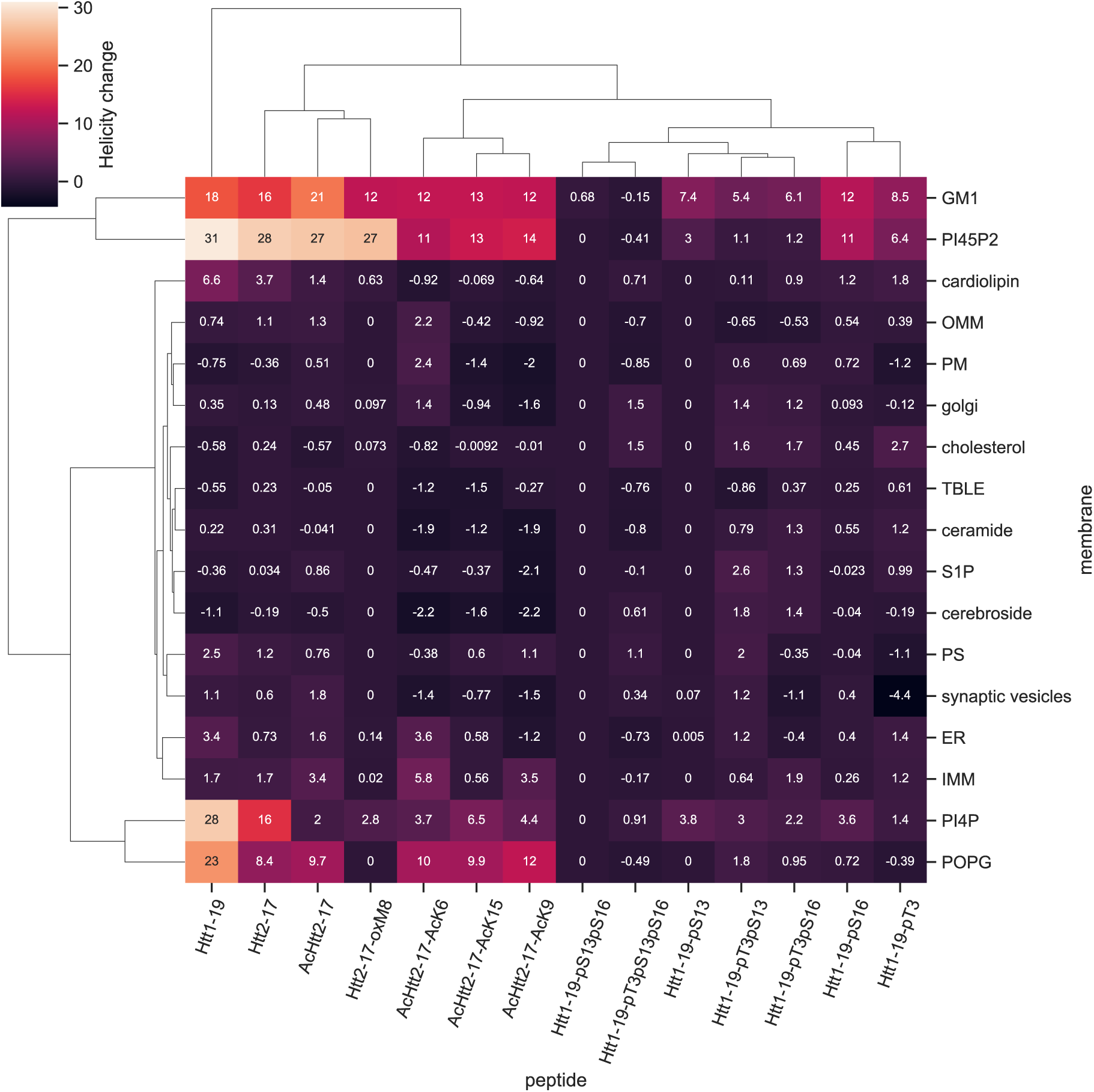
Lipid-dependent helical shift of Httex1 N-terminal peptides. The helicities (%) of unmodified and PTM’ed Nt17/19 were measured by CD in the presence and absence of lipids. The helical changes were calculated by subtracting the in-solution helicity of Nt17/19 from its helicity in the presence of liposomes made of 100 % POPG, liposomes made of brain lipid extract spiked with 50% mol of specific lipids, or mimicking the lipid composition of organelle membranes. Hierarchical clustering was performed on the heatmap for both peptides and membranes, highlighting the PTMs and lipids that cause similar effects on helicity change.

We observed that in general, the presence of membranes increases the helicity of both peptides, as discerned by CD (Figure 1, Figure S1). Previously, we showed that unmodified Nt17/19 formed an α-helix rich conformation upon incubation with vesicles made of 100% mol phosphatidylglycerol (POPG) at 5 molar equivalent ^14^. Herein, our data show that membranes made of at least 50% mol of specific anionic lipids, such as the phosphoinositide PI4P, PI(4,5)P2, and the ganglioside GM1 strongly increased the helical content of unmodified Nt17/19 (Figure 1). To a lesser extent, membrane models containing 50% mol cardiolipin (CL) or mimicking organelle with a low packing environment like endoplasmic reticulum (ER) and inner mitochondrial membrane (IMM) mimics also increased the percentage of α-helical contents of Nt17/19 WT (Figure 1). Similar to the unmodified Nt17/19, all Nt17 peptides with single acetylation at K6, K9, or K15 exhibited increased helicity in the presence of POPG, PI(4,5)P2, GM1, but this increase in helicity was lower in these single acetylated peptides compared to the unmodified Nt17 (Figure 1). We also examined the effect of phosphorylation on Nt19 membrane interaction. Single serine phosphorylation at residue 13 and 16 decreased but did not abolish the POPG and PIP2-induced helicity. Phosphorylation at T3 increased helicity in a membrane-independent manner, whereas phosphorylation of both serine residues (pS13 and pS16) significantly reduced Nt17 helicity regardless of the lipid composition (Figure 1, Figure S2). Next, we studied the effects of M8 oxidation on Nt17 conformation in the presence of membranes. OxM8 decreased membrane-induced helicity in a lipid-dependent manner. oxM8-Nt17 becomes helical in the presence of PI(4,5)P2 and GM1, and to a lesser extent in the presence of PI4P. No change in oxM8 conformation was observed in the presence 100% POPG and 50% cardiolipin, unlike the unmodified Nt17 (Figure 1).

These results demonstrate that lipid composition and PTMs could play important roles in differentially regulating helical conformation of Nt17/19 on membranes. Overall, GM1 and PI45P2 had the most significant impact on increasing the helicity of most PTM’ed Nt19. PI4P and POPG behaved similarly to GM1 and PI45P2 in increasing helicity. Interestingly, these two lipids showed much less enhancement of helicity for Nt17 with oxM8. Cardiolipin only increased helicity for unmodified Nt17/19 but had no significant effects on the acetylated, oxidized, or phosphorylated peptides. Meanwhile, cholesterol, ceramide, S1P, cerebroside, and PS had no significant impact on the helicity of any of the unmodified and modified Nt17/19 peptides. These results demonstrate that PTMs exert different roles in different membrane environments, thus highlighting how PTMs could increase the functional and structural diversity of proteins.

### Molecular mechanism of PTMs’ impact on Nt19 membrane interaction

To better understand how membrane composition influences the conformational properties of Nt17 and how PTMs influence the interaction between Nt17/19 and membranes, we conducted MD studies on differentially modified Nt19 in the presence of 100% POPG membrane, plasma membrane mimetics, and synaptic vesicle mimetics. The lipid compositions of physiological mimetics are the same as the ones used in our *in vitro* CD studies. The simulations were run with CHARMM36m force field and a modified TIP3P water model ^51,52^.

We first performed clustering with CLoNe ^53^ based on backbone dihedral angles for the 13 µs simulation for Nt19 corresponding in-solution simulations (Figure S3) ^15^. The major conformations (Figure S4) were then used as starting conformations for 5 µs simulation in the presence of a membrane and placed 46 Å from the center of the bilayer.

First, we conducted simulations for unmodified, pT3, AcK6, and AcK9 Nt19 in the presence of POPG (Figure 2). The major conformations we extracted from WT Nt19 in solution are a short C-term helix (C1), longer N-term helix (C2), and a disordered conformation (C3) (Figure S4). In the presence of POPG membranes, the WT C2 conformation, found with high abundance (34.5%) in in-solution MD simulation, maintained its helicity throughout the simulation, while the shorter C-term helical WT C1 (4.5% abundance in solution) and disordered WT C3 conformations (61.0% abundance in solution) only formed unstable and very short helices. For AcK6-Nt19, the high abundance AcK6 C1 conformation, which is a long helix, formed relatively stable helices on the membrane during the simulation. For AcK9-Nt19, the abundant disordered conformation, AcK9-Nt19 C4, folded into a helix and remained helical throughout the simulation. The low abundancy short N-term helix AcK9-Nt19 C1 remained similar conformation on the membrane. These findings are consistent with the experimental result demonstrating that unmodified and acetylated Nt19 exhibit increased helicity in the presence of POPG membranes and showed that this increased helicity might result from the stabilization of helical conformations on the membrane or the facilitated disordered-helical conformation transition on the membrane.

**Figure 2.**
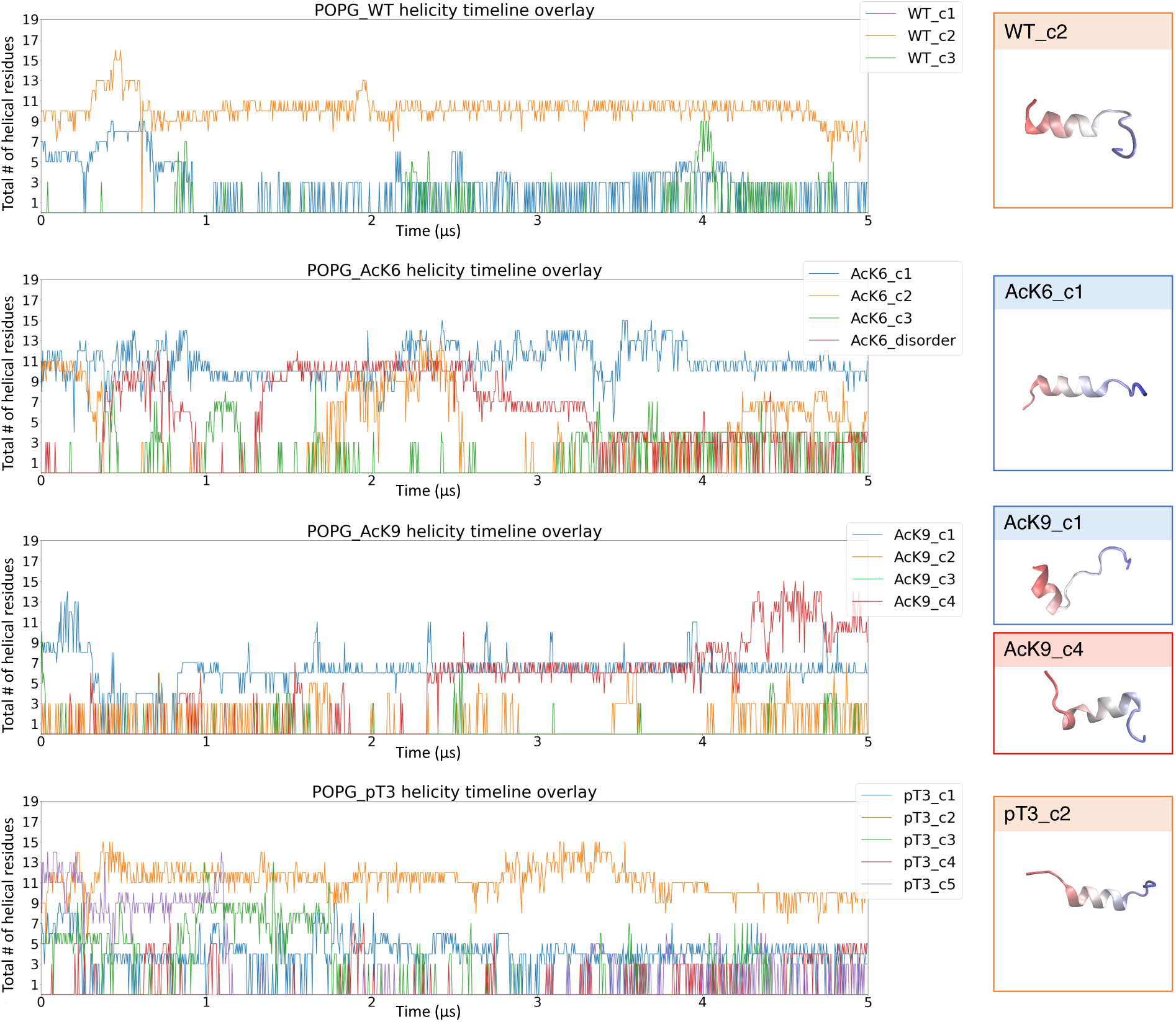
Certain in-solution conformations were particularly stabilized by the membrane. The total number of helical residues was obtained from molecular simulations for the whole trajectory. The conformations stabilized on the membrane after 5 µs were shown on the right. Nt19 WT c2, AcK6 c1, AcK9 c1/c4, pT3 c2 helicity was stabilized by POPG membrane. The molecules are colored red to blue from N-term to C-term.

Although the unmodified and acetylated Nt19 peptides exhibit increased helicity in the presence of certain membranes (as assessed by CD), pT3-Nt19’s helicity is not sensitive to the presence of any membranes, irrespective of the lipid compositions (Figure 1). This could be due to reduced interaction with the membrane or the fact that pT3-Nt19 already forms a stable helix in solution on its own ^9^. Simulation for unmodified Nt19 helical major conformation C1 showed a rapid formation of a stable helix on the membrane within the first 50 ns. In contrast, the major helical conformation of pT3-Nt19 either has an unstable contact with the membrane (conformation C3) or forms stable contact with the membrane only after unfolding

(conformation C5) (Figure 2, Figure 3). This is likely due to the negatively charged phosphate group, which disfavors the interaction with the negatively charged lipid groups. Interestingly, when we started the simulation with a disordered conformation of pT3, it seems to have similar contact with the POPG membrane as the disordered conformation of unmodified Nt19 (Figure S5), indicating that the flexibility of fully disordered conformation might allow Nt19 to adapt conformations that lower the impact of phosphorylation on membrane interaction. However, despite the binding of disordered pT3-Nt19 to the membrane, it remained in disordered conformation rather than having enhanced helicity by the membrane. Thus, the insensitivity of pT3’s helicity to the presence of membranes observed in experiments was likely caused by both the lowered interaction with the membrane and the lack of changes in its helicity even when binding to the membrane.

**Figure 3.**
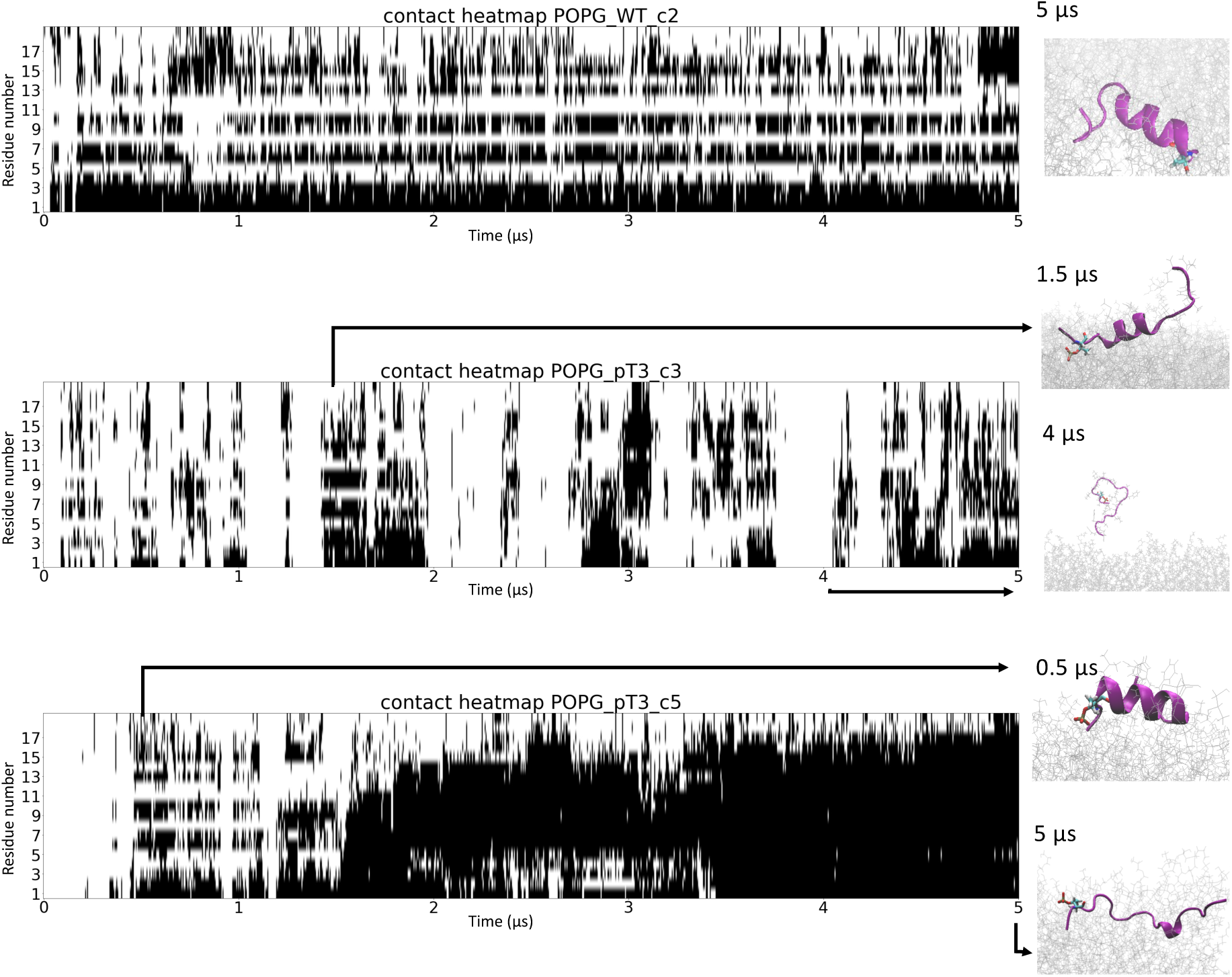
Phosphorylation at T3 disrupted the stabilization of Nt19 helical conformation by the membrane. The distance between each residue and the lipid membrane was calculated throughout the molecular simulation trajectory. The residues within 6 Å are marked as black. The conformations corresponding to the marked time points are shown on the right. These contact heatmaps indicated that the helical conformation of unmodified Nt19 formed stable periodic contact with the membrane, while the pT3 helical conformation had reduced and unstable contact with the membrane.

Our results demonstrate the utility of using atomistic simulation for studying Nt17-membrane interaction. While coarse-grained simulations are frequently favored for studying protein-membrane interactions due to the longer time scale they can access and reduced computational demands, they fall short when modeling Nt17-membrane interaction. As a highly dynamic peptide, Nt17 undergoes complex unfolding and refolding processes when interacting with membranes. These processes involve various conformations: some maintain stable interaction with the membrane and remain the same fold on the membrane, others interact with the membrane but unfold into disordered states or refold into novel conformations, while some do not stably adhere to the membrane. Given this complexity, coarse-grained simulations simply cannot realistically capture the complex interaction between Nt17 and membrane. Atomistic simulations are capable of capturing the dynamics of folding and unfolding of Nt17 on the membrane, offering precise positioning and orientation of amino acid side chains, and providing more accurate representation of interactions. All of which are essential for understanding PTM-mediated Nt17-membrane interactions.

### Lysine residues are essential for membrane interaction

The Nt19 with single acetylation at K6, K9, and K15 showed increased helicity, as discerned by CD spectroscopy (Figure 1) in the presence of certain negatively charged membranes, even though acetylation at all three lysine residues was shown to decrease Nt17 membrane interaction ^26^. In our simulation, residue-wise contact of unmodified Nt19 showed a stable periodic pattern for the most abundant helical conformation in-solution, and the lysine residues showed extensive contact with the membrane (Figure 4A). However, simulations of major conformations of AcK6 showed that certain helical conformations still formed stable contact with the membrane and maintained stable conformation on the membrane (Figure 4B). Thus, we further investigated the importance of lysine residues for Nt19-membranes interaction and the impact of single and multiple lysine acetylation on such interaction. We conducted simulations with the helical conformation C2 of unmodified Nt19 and also simulations of this conformation with mono- (AcK6-Nt19), bi- (Ack6/AcK9-Nt19), and tri-acetylation (Ack6/AcK9/AcK15-Nt19) in the presence of POPG membrane. Simulation results showed that single acetylation does not influence membrane interaction of Nt19, while bi- and tri-acetylation nearly abolished the membrane interaction (Figure 4C). This result demonstrated the possible regulation of HTT-membrane interaction by the acetylation level of Nt17/19.

**Figure 4.**
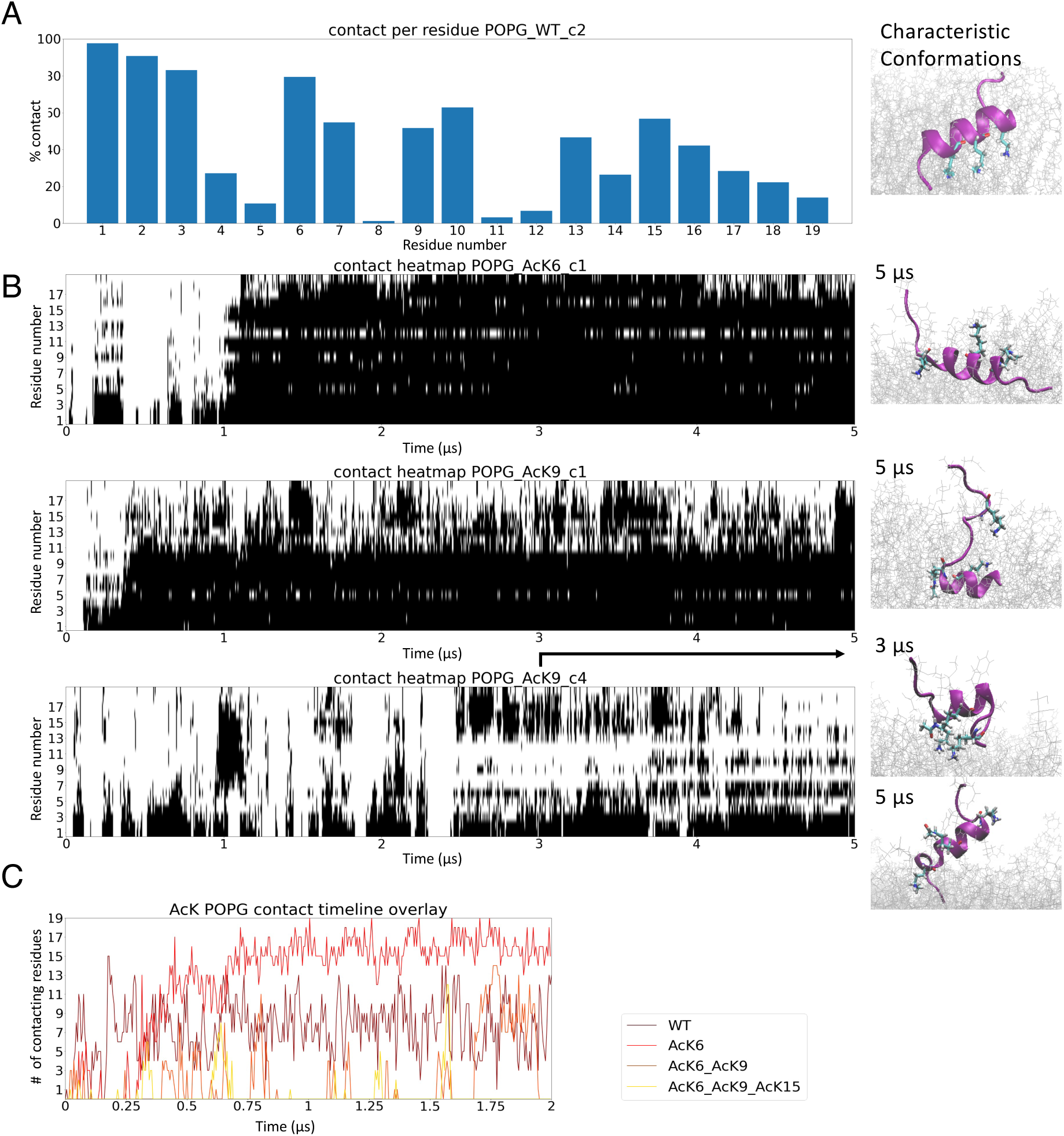
Effect of acetylated lysine for Nt19 membrane interaction. (A) The contact with the membrane of each residue was averaged over the whole trajectory of unmodified Nt19. This showed that lysine residues have particularly high interaction with lipids. (B) Contact heatmap for Nt19 with lysine acetylation from MD showed that single lysine acetylation was not enough to abolish Nt19-membrane interaction. The conformations corresponding to the indicated time points are shown on the right. (C) Overlay of total number of residues contacting membrane from MD for unmodified, mono-, di-, and tri-acetylated Nt19 demonstrated that di- and tri-acetylated Nt19 had much lower interaction with POPG compared to the unmodified Nt19, while mono-acetylated Nt19 had membrane interaction similar to the unmodified Nt19.

### Enrichment of SV mimetics with PI45P2 resulted in increased stabilization of helical conformations

In addition to POPG, lipids like PI4P, PI45P2, GM1, and cardiolipin were also shown to increase the helicity of Nt19 in our CD experiments; thus, to understand the molecular basis of the influence of these lipids on HTT-membrane interaction, we simulated Nt19 in the presence of SV mimetics and plasma membrane mimetics.

We first conducted simulations with SV enriched with 50% PI45P2, since it has critical roles in the nervous system. Simulations were conducted with the two high abundance conformations of unmodified Nt19, one partially helical (C2) and the other one disordered (C3). We compared the helical time evolution for these two conformations in the presence of SV mimetics and in the presence of plasma mimetics. As shown in Figure 5A, the membrane enriched with PI45P2, the unmodified Nt19 C2 helical conformation remained relatively stable, while the C3 disordered conformation more rapidly folded into a helical conformation. Also, the C2 helical conformation seems to preferentially contact the membrane with its lysine residues in the presence of a membrane enriched with PI45P2. Further, we have noticed that this significant interaction between Nt19 and PI45P2 could be achieved with ionic interactions between the lysine groups of Nt19 and the phosphate group of PI45P2 (Figure 5B). The extensive interaction between lysine residues and the membrane was also observed in multiple unmodified and modified Nt19 conformations that stabilized helices on the membrane.

**Figure 5.**
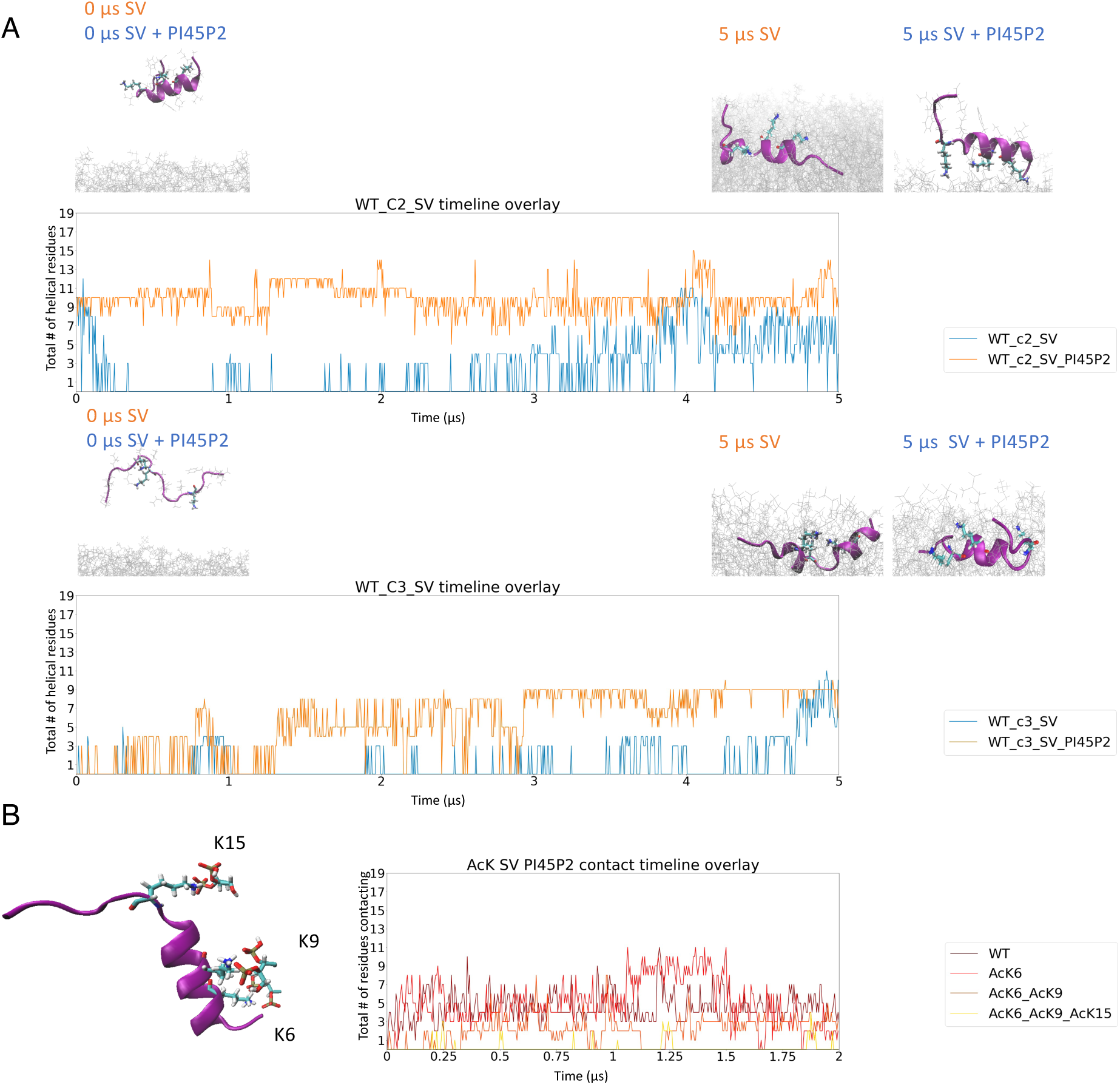
Nt19 WT preserves helical conformation better on SV enriched with PI45P2. (A) Overlay of total number of helical residues from MD for unmodified Nt19 in the presence of SV mimetics and SV mimetics enriched with 50% PI45P2 showed that PI45P2 stabilized the helical conformation (WT c2) and induced faster folding into helical conformation from disordered conformation (WT c3). The conformations at the start and end of the MD are shown above the plot. (B) A snapshot from the simulation of unmodified Nt19 in the presence of SV mimetics (left) and the total number of residues contacting the membrane throughout the simulation (right) showed that the salt bridge between phosphate of lipid and lysine stabilizes the interaction between Nt19 and membrane. Di- and tri-acetylation of Nt19 resulted in decreased interaction with SV mimetics enriched with PI45P2.

This prompted us to further investigate the role of positively charged lysine residues in the interaction with SV membrane enriched with PI45P2. Since the positively charged lysine residues seem to form extensive contact with certain negatively charged lipids, we hypothesized that the interaction between Nt19 and membrane could be suppressed by acetylating these lysine residues. Thus, we simulated differentially acetylated Nt19 in the presence of synaptic vesicle mimetics enriched with 50% PI45P2. With 2 µs MD simulations, we observed that the interaction between membranes and Nt19 with AcK6/AcK9 or AcK6/AcK9/AcK15 was lower than the single acetylated and unmodified Nt19 (Figure 5B). This supported our hypothesis that the positively charged lysine residues have important roles in the regulation of the interaction between Nt19 and membranes.

Finally, we conducted a simulation with plasma membrane mimetics, which contains 5% PI45P2, and analyzed the normalized contacts between each lipid type in the plasma membrane with Nt19. We conducted the analysis with both the helical in-solution conformation of Nt19 (C2) and the disordered in-solution conformation of Nt19 (C3). Throughout the entire simulation of both, PI45P2 was shown to have more contact with Nt19 compared to other lipids (Figure 6A). Furthermore, volumetric maps for the occupancy of each lipid averaged over the whole trajectory also showed that protein got centered around by PI45P2 (Figure 6B), even though the lipids were randomly placed in the beginning of the simulations. This result demonstrated that lipids like PI45P2 might play important roles in driving Nt19-membrane interactions despite their low abundance in the physiological membranes.

**Figure 6.**
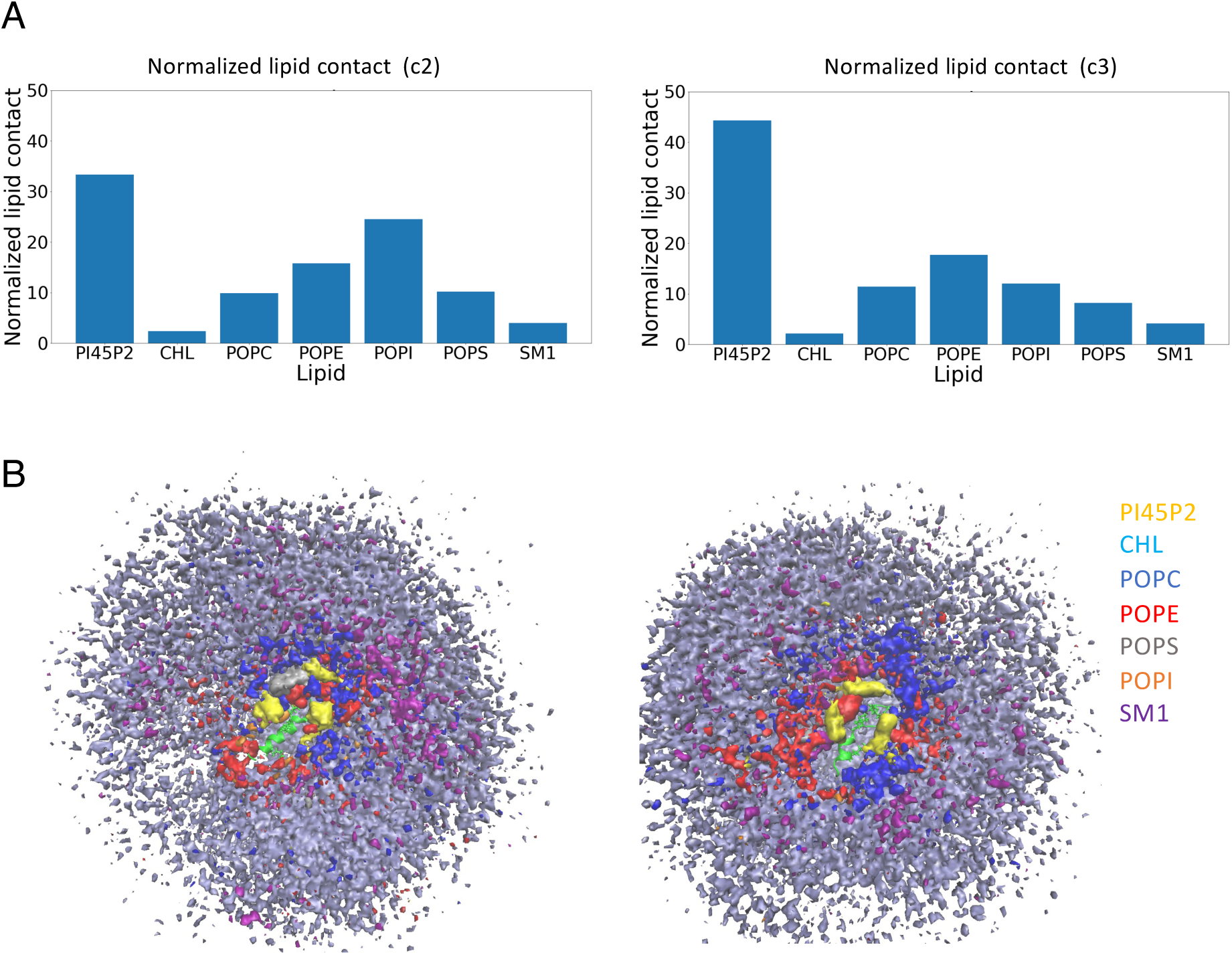
PI45P2 has a particularly high interaction with unmodified Nt19 compared to other lipids. (A) Contact of each type of lipid in the plasma membrane throughout the trajectory and normalized by the abundance of the lipid. The analysis was done for both the helical (left) and the disordered in-solution conformation of unmodified Nt19 (right) as starting conformation, and PI45P2 has more extensive contact with Nt19 compared to other lipids in both cases. (B) Occupancy of each lipid type averaged over the whole trajectory obtained by in VMD. PI45P2 gathered around Nt19 (in green) even though the lipids were randomly distributed at the beginning of the simulations.

The significance of the finding is underscored by the pivotal role PI45P2 plays in the nervous system, particularly in regulating various facets of the synaptic vesicle cycle. Supporting this, disruptions in phosphoinositides levels and signaling have been identified in numerous dementias ^54^. Notably, both Alzheimer’s Disease (AD) and Parkinson’s Disease (PD) manifest alterations in the expression of phosphoinositide conversion enzymes, pinpointing these enzymes as promising therapeutic targets ^55,56^.

### Conclusions

Our results highlight the inherent complexity and intricate differential effects of PTMs on HTT-membrane interaction dynamics. These findings underscore the necessity for a deeper understanding of these interactions in the presence of PTM crosstalks, as investigating PTMs one at a time could mask important insight into the molecular determinants of HTT interactions with membranes in health and disease. Further studies are necessary to understand how the complex interplay between PTMs, Nt17 conformation, and membrane composition influence the structure and aggregation mechanisms of mutant Httex1 or other longer N-terminal Htt fragments. Insights from these studies could pave the way for a better understanding of how changes in lipids and the dynamic properties of PTMs could trigger or alter HTT aggregation and toxicity, facilitating the discovery of novel therapeutic strategies for the treatment of HD.

## Materials and Methods

### Materials and Instrumentation

The peptides used in this work were synthesized by Integrated Research Biotech Model (IRBM) through a collaboration with the CHDI Foundation. The lipids DOPC, POPC, DOPE, POPE, DOPI, POPI, DOPS, POPS, Brain PS, POPG, Total Brain lipid extract, Brain SM, Brain GM1, Brain cerebrosides, Brain ceramides, cardiolipin, sphingosine, sphingosine-1-phosphate, cholesterol, Brain PI4P, Brain PI(4,5)P_2_ and Brain PS were purchased from Avanti Polar Lipids. HPLC-grade acetonitrile was purchased from Macherey-Nagel. Formvar carbon film on 200-mesh copper grids (FCF200-Cu) and uranyl formate (16984-59-1) from Electron Microscopy Sciences were used for sample preparation for negative-stain transmission electron microscopy (TEM).

### Nt17 peptide analysis

Each peptide was analyzed for purity by LC-ESI-MS and C8-UPLC (Figure S6). Liquid chromatography electrospray ionization mass spectrometry (LC-ESI-MS) was performed using a C3 column (Agilent Poroshell 300SB-C3, 1 x 75 mm, 5 µm) with a gradient of 5% to 95% acetonitrile (0.1% v/v formic acid) at 0.3 mL/min over 10 minutes with UV detection at 214 nm and MS detection on a Thermo LTQ. Mass spectra were deconvoluted using the MagTran v. 1.03b software (Amgen). Ultra performance liquid chromatography (UPLC) analysis was done on a Waters UPLC system (Waters Acquity H-Class) using a C8 column (Waters Acquity UPLC BEH300, 1.7 μm, 300Å, 2.1x150 mm) with UV detection at 214 nm using a 2.75 minute gradient of 10% to 90% acetonitrile (0.1% v/v trifluoroacetic acid) at 0.6 mL/min. Microcuvettes for density light scattering were purchased from Wyatt Technology.

### Vesicle formation

Vesicles were prepared using the extrusion method. Briefly, individual phospholipids or phospholipid mixtures in chloroform were dried using a nitrogen stream to form a thin film on the wall of a glass vial. Potentially remaining chloroform was removed by placing the vial under vacuum overnight. The phospholipids were solubilized in PBS to the desired final concentration by sonication. The solution was then passed through an Avestin LiposoFast extruder (Avestin Inc.) (membrane pore size: 0.1 µM), and the size and homogeneity of the resulting vesicles were assessed by DLS and EM.

### Density Light Scattering

The size distribution of prepared vesicles (Figure S7A) was analyzed by density light scattering (DLS) in microcuvettes using a DynaPro NanoStar from Wyatt Technology.

### Transmission electron microscopy

For TEM analysis (Figure S7B), 3 µl of liposomes solution was spotted onto a Formvar/carbon-coated 200-mesh glow-discharged copper grid for 1 min. The grid was then washed twice times with water, once with 0.7% (w/v) uranyl formate and stained for 30 seconds with 0.7% w/v uranyl formate. Imaging was performed on a Tecnai Spirit BioTWIN electron microscope equipped with a LaB6 gun and a 4K x 4K FEI Eagle CCD camera (FEI) and operated at 80kV.

### Membrane binding assay

Samples for membrane binding studies were prepared by adding 5 molar equivalents of lipids (300 µM, final concentration) to the Nt17 peptides solution (60 µM, final concentration) and incubating the peptide/LUV samples at RT for 1 hour before CD measurements.

### Circular dichroism spectroscopy

For CD analysis of the Nt17 peptides, 10-200µM of proteins were prepared in 200µL filtered PBS and analyzed using a Chirascan Plus CD spectrometer from Aimil in a 1-mm quartz cuvette. The ellipticity was measured from 195 to 250 nm at 25 °C, and the data points were acquired continuously every 0.5 nm and a bandwidth of 1.0 nm, 0.5s per point. Three spectra of each condition were obtained and averaged. A sample containing buffer only was analyzed and subtracted from each signal. The obtained spectra were smoothed using a binomial filter, order 2, and the resulting spectra were plotted as the mean residue molar ellipticity (θMRE).

### Peptides concentration dependent studies

For peptide concentration dependent studies, each peptide was weighed in a separate Eppendorf tube and dissolved in filtered phosphate buffered saline (0.45µm) at the desired concentration.

### pH dependent studies

For pH dependence studies, each peptide was weighed in a separate Eppendorf tube for each condition and dissolved in filtered buffer (0.45µm) at the desired pH (pH 3, 4 and 5, 10 mM Sodium acetate, 75 mM NaCl; pH 6, 7 and 8, 10 mM Sodium phosphate, 75mM NaCl; pH 9-10, Sodium carbonate 10mM, 75 mM NaCl). The secondary structure of each sample was then measured by CD. To determine the helical content at each pH we used the web server BestSel.

### Preparation of Starting conformation for Nt19-membrane simulations

In our previous study ^15^, we conducted 13 μs simulations for unmodified, phosphorylated, and acetylated Nt19 in solution with explicit CHARMM36m ^52^ and modified TIP3P water model.

We calculated dihedral angles for the Nt19 backbone, and a transformation from the space of dihedral angles (ϕ_n_, ψ_n_) to metric coordinate space was done by taking trigonometric function (cosϕ_n_, sinϕ_n_, cosψ_n_, sinψ_n_). Then, we performed dihedral angle principle component analysis (dPCA) ^57^, clustering based on the first 3 components, and major conformation extraction with CloNe ^53^.

### Molecular dynamics simulations

The starting structures of Nt19 were the major conformations extracted from 13 µs in-solution simulations with CLoNe ^53^ based on backbone dihedral angles. The inputs were generated with CHARMM-GUI Membrane Builder ^58^, and the size of the membrane surface was 80 Å * 80 Å. The protein was placed 46 Å from the center of the bilayer. The first principal axis was aligned along the Z axis of the system, and the peptide was rotated with respect to the X axis for 90 degree. A water layer of thickness 50 Å was added to the system. The solvation was done with explicit CHARMM36m ^52^ modified TIP3P water model and 0.14 M Na^+^ and Cl^-^ ions. The simulations were conducted with GROMACS ^59^. CHARMM36m forcefield with modified residues was used for phosphorylated threonine and acetylated lysine residues ^60^, and the phosphate group of phosphorylated threonine was in a dianionic form. The system was minimized with steepest descent and equilibrated to 1 atm and 303.15 K through 6 cycles with slowly reducing restraint forces. The nonbonded interaction cut-off was 12 Å. The time step was 1 fs for equilibration and 2 fs for production. The Berendsen temperature coupling method was used to maintain temperature, and the semi-isotropic Berendsen method was used for pressure coupling ^61^. LINCS algorithm ^62^ was used to constrain the bond distances. The production trajectories were run for 2 μs for simulations of differentially acetylated Nt19 and 5 μs for all other systems.

## Acknowledgements

We would like to thank Syed Muhammed Muazzam Kamil for peptide sample preparation. We would like to thank Juan Francisco Bada Juarez for the valuable discussion for article preparation. We would like to thank all members of Laboratory of Molecular and Chemical Biology of Neurodegeneration and Laboratory for Biomolecular Modeling for their feedback and input on the project.

## Author Contributions

HL and MDP supervised all the work, CG and ZZ performed the experiments, ZZ and LA conducted and analyzed molecular simulation results, CG and ZZ prepared the first draft of the paper and all authors contributed to the writing and preparation of the final version.

