## supplemental file for "Differential Effects of Posttranslational Modifications on the Membrane Interaction of Huntingtin Protein"

### Supplementary material

| Peptides | Sequence | Expected Mass (Da) | Isoelectric point |
|---|---|---|---|
| Htt2-17 | ATLEKLMKAFESLKSF | 1843 | 9.8 |
| Htt2-17 OxM8 | ATLEKL**(Mox)**KAFESLKSF | 1859 | |
| AcHtt2-17 | **(Ac)**ATLEKLMKAFESLKSF | 1885 | 7.32 |
| Ac-Htt2-17-AcK6 | **(Ac)**ATLEK**(Ac)**LMKAFESLKSF | 1927 | 4.15 |
| Ac-Htt2-17-AcK9 | **(Ac)**ATLEKLMK**(Ac)**AFESLKSF | 1927 | 4.15 |
| Ac-Htt2-17-AcK15 | **(Ac)**ATLEKLMKAFESLKSF**(Ac)** | 1927 | 4.15 |
| Htt1-19 | MATLEKLMKAFESLKSFQQ | 2230 | |
| Htt1-19-pT3 | MA**(p)**TLEKLMKAFESLKSFQQ | 2310 | |
| Htt1-19-pS13 | MATLEKLMKAFE**(p)**SLKSFQQ | 2310 | |
| Htt1-19-pS16 | MATLEKLMKAFESLK**(p)**SFQQ | 2310 | |
| Htt1-19-pT3/S13 | MA**(p)**TLEKLMKAFE**(p)**SLKSFQQ | 2390 | |
| Htt1-19-pT3/S16 | MA**(p)**TLEKLMKAFESLK**(p)**SFQQ | 2390 | |
| Htt1-19-pS13/S16 | MATLEKLMKAFE**(p)**SLK**(p)**SFQQ | 2390 | |
| Htt1-19-pT3/S13/S16 | MA**(p)**TLEKLMKAFE**(p)**SLK**(p)**SFQQ | 2470 | |

**Table S1. Sequence, composition and biochemical properties of peptides used in liposome binding assay.** Ac: acetylation, p: phosphorylation, Ox:oxidation

| class | lipid | shape | Net charge at pH7 | Cell localization | %mol in cells | Link with Htt |
|---|---|---|---|---|---|---|
| **mix** | **Brain lipid extract (TBLE)** | mix | 0 | mix | 100% | HD is a Neurodegenerative disease |
| **sphingolipids** | **Sphingosine-1-phosphate (S1P)** | Inverted cone | -1 | ER, PM, mitochondria | <1% | (Di Pardo and Maglione, 2018) |
| | **ceramides** | Cylindrical | 0 | ER, PM, Golgi, lysosomes, mitochondria | <5% | Precursor of cerebrosides, SM and GM1 |
| | **cerebrosides** | Cylindrical | 0 | Golgi, PM | <1% | Precursor of GM1 |
| | **Ganglioside GM1** | Inverted cone | negative | PM outer leaflet, lysosomes | 6% in neurons | (Chaibva et al., 2018) |
| **glycerophospholipids** | **cardiolipin** | Cylindrical | -2 | Mitochondria | 5-20% in mitochondria | (Kegel et al., 2009) |
| | **Phosphoinositide-4-phosphate (PI4P)** | Inverted cone | -3 | Golgi | <1% | (Kegel et al., 2009) |
| | **Phosphoinositide-4,5-biphosphate ($PI(4,5)P_2$)** | Inverted cone | -4 | PM inner leaflet | <1% | (Kegel et al., 2009) |
| | **Phosphatidylglycerol (PG)** | cylindrical | -1 | mitochondria | <5% | (Chiki et al., 2017; DeGuire et al., 2018) |
| | **Phosphatidylserine (PS)** | cylindrical | -1 | All organelles | <10% | (Kegel et al., 2009; Michalek et al., 2013) |
| **sterols** | **cholesterol** | Inserted inside the bilayer | 0 | All organelles | 40% | (Gao et al., 2016; Michalek et al., 2013) |

**Table S2. Biophysical properties and biological significance of tested lipids.**

| Lipid class | Studied lipid | liposomes | Lipid composition (%mol) |
|---|---|---|---|
| mix | Brain lipid extract (TBLE) | TBLE | Brain total lipid extract (100%) |
| sphingolipids | Sphingosine-1-phosphate | S1P | TBLE (50%); Brain sphingosine-1-phosphate (50%) |
| | ceramides | ceramides | TBLE (50%); Brain ceramides (50%) |
| | cerebrosides | cerebrosides | TBLE (50%); Brain cerebrosides (50%) |
| | Ganglioside GM1 | GM1 | TBLE (50%); Brain ganglioside GM1 (50%) |
| glycerophospholipids | cardiolipin | cardiolipin | TBLE (50%); 18:1/18:1 cardiolipin (50%) |
| | Phosphoinositide-4-phosphate (PI4P) | PI4P | TBLE (50%); Brain PI4P (50%) |
| | Phosphoinositide-4,5-biphosphate (PI(4,5)$P_2$) | PI(4,5)$P_2$ | TBLE (50%); Brain PI(4,5)$P_2$ (50%) |
| | Phosphatidylglycerol (PG) | POPG | POPG (100%) |
| | Phosphatidylserine (PS) | PS | TBLE (50%); Brain PS (50%) |
| sterols | cholesterol | cholesterol | TBLE (50%); cholesterol (50%) |

**Table S3. Lipid composition of simple liposomes.**

| Organelle surrogate | Liposomes | Lipid composition (%mol) |
|---|---|---|
| **Endoplasmic reticulum** (van Meer et al., 2008) | ER | DOPC (60%), DOPE (20%), DOPI (10%), DOPS (5%), cholesterol (5%) |
| **Mitochondria** (van Meer et al., 2008; Valencak and Azzu, 2014) | IMM | DOPC (40%), DOPE (40%), cardiolipin (15%), DOPI (5%) |
| | OMM | DOPC (50%), DOPE (30%), DOPI (10%), DOPS (5%), cardiolipin (5%) |
| **Golgi apparatus** (Bigay et al., 2003) | Golgi | POPC (50%), POPE (19%), cholesterol (16%), POPI (10%), POPS (5%) |
| **Plasma membrane** (Temmerman and Nickel, 2009) | PM | Cholesterol (50%), POPC (12.5%), Brain SM (12.5%), POPE (10%), POPI (5%), POPS (5%) |
| **Synaptic vesicles** (Fusco et al., 2014) | SV | 50% DOPE (50%), DOPS (30%), DOPC (20%) |

**Table S4. Lipid composition of liposomes mimicking the lipid composition of organelle membranes.**

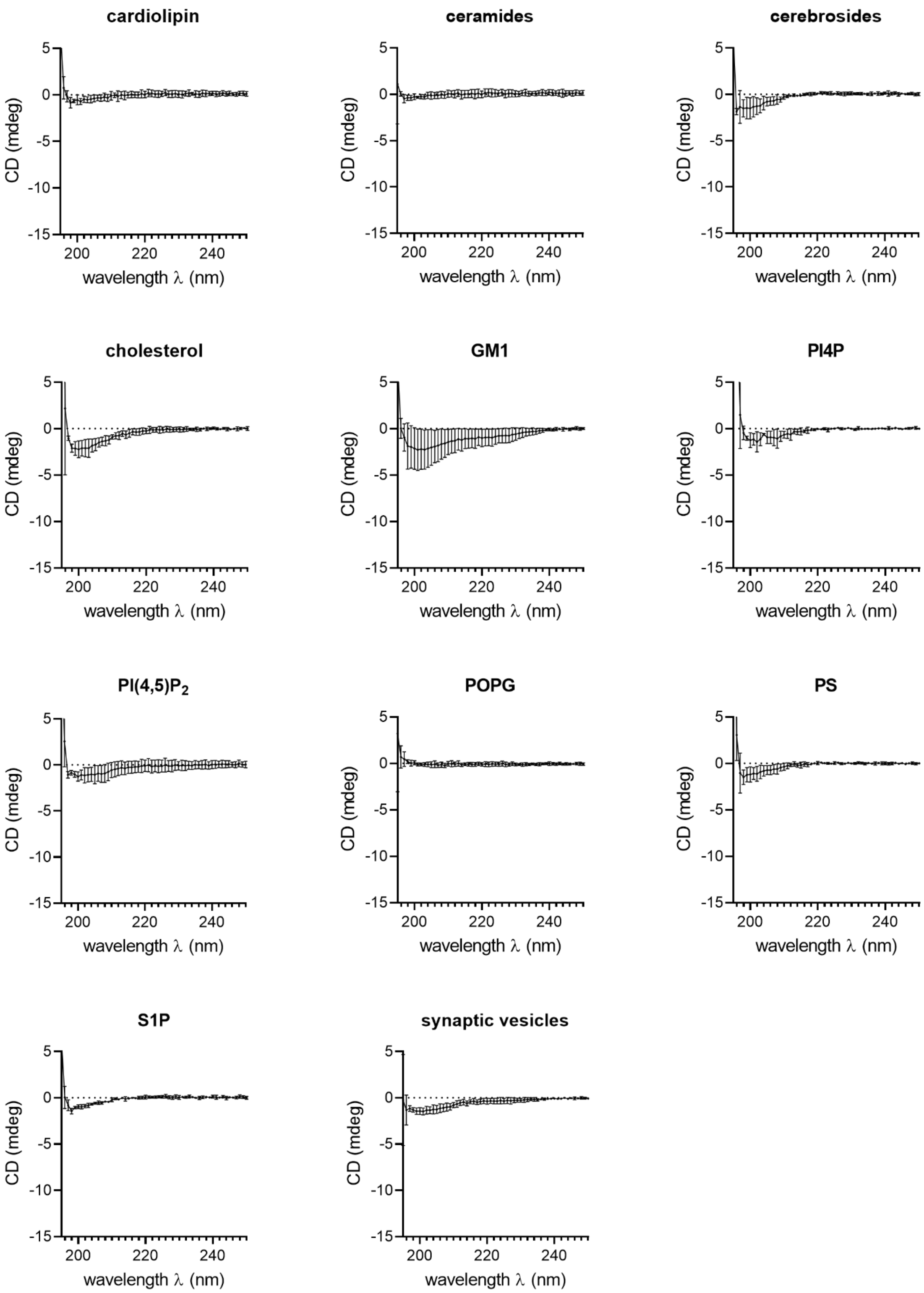
cardiolipin
ceramides
cerebrosides
cholesterol
GM1
PI4P
PI(4,5)P2
POPG
PS
S1P
synaptic vesicles
CD (mdeg)
wavelength λ (nm)
5
0
-5
-10
-15
200
220
240

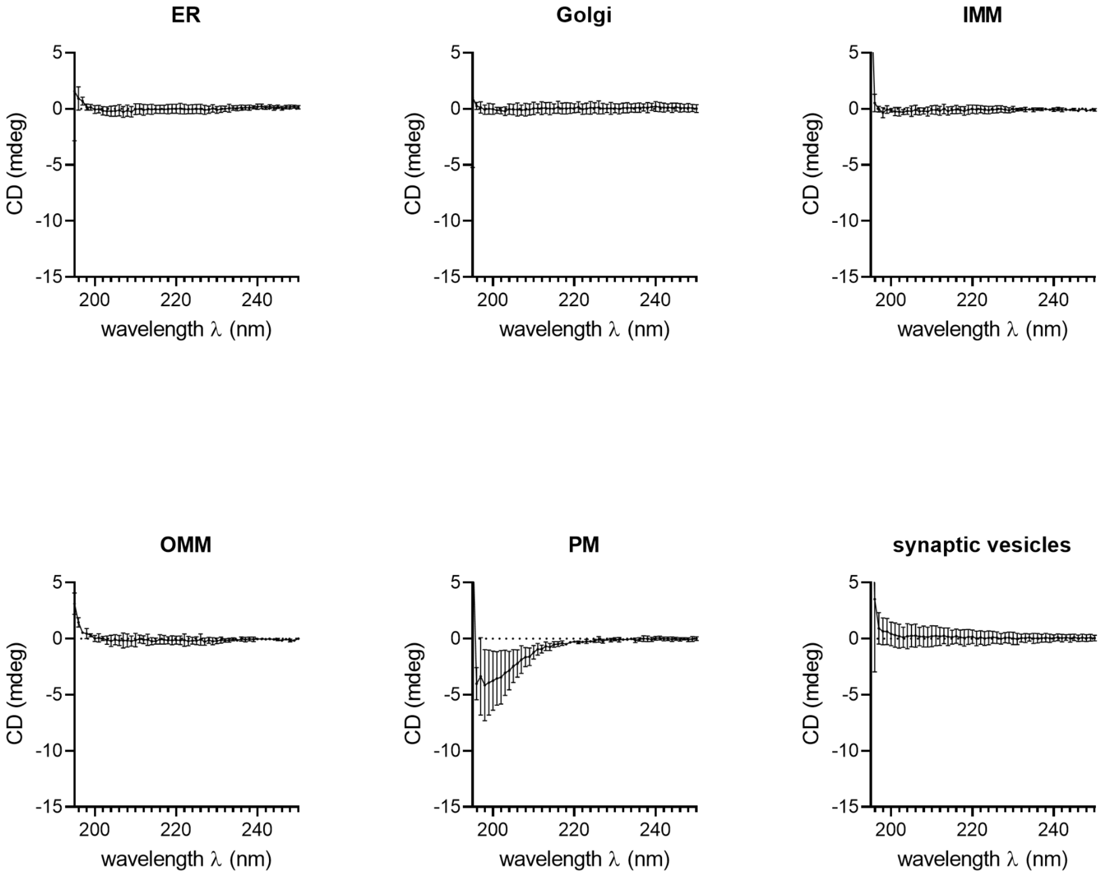


**Figure S1. CD spectra of the liposome-containing buffers.**

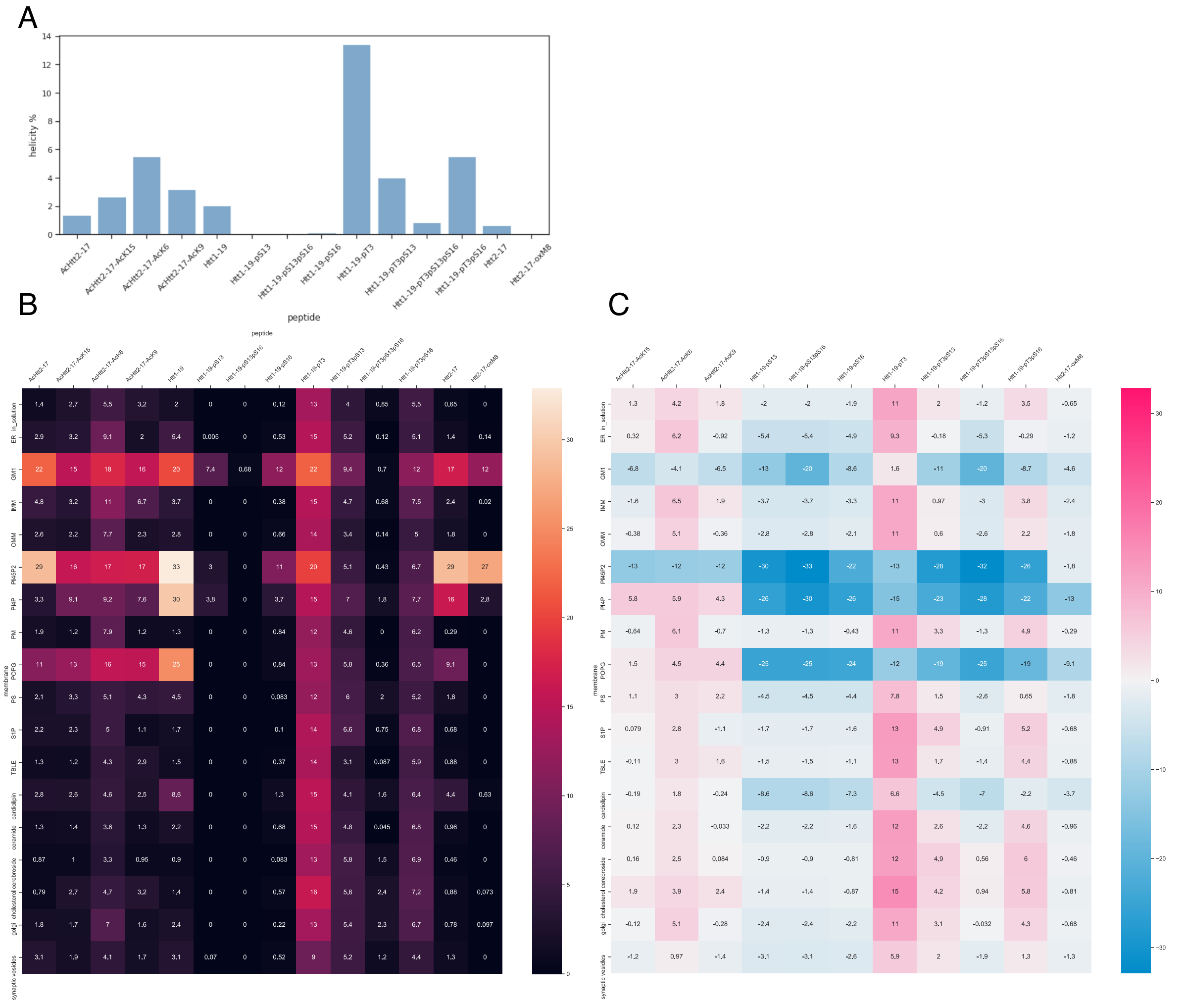

**Figure S2. (A) Helicity of unmodified, acetylated, and phosphorylated Nt17/19 in solution. (B) Averaged helicity of Nt17/19 in the presence of different membranes. (C) Helicity changes resulted from the addition of PTMs.**

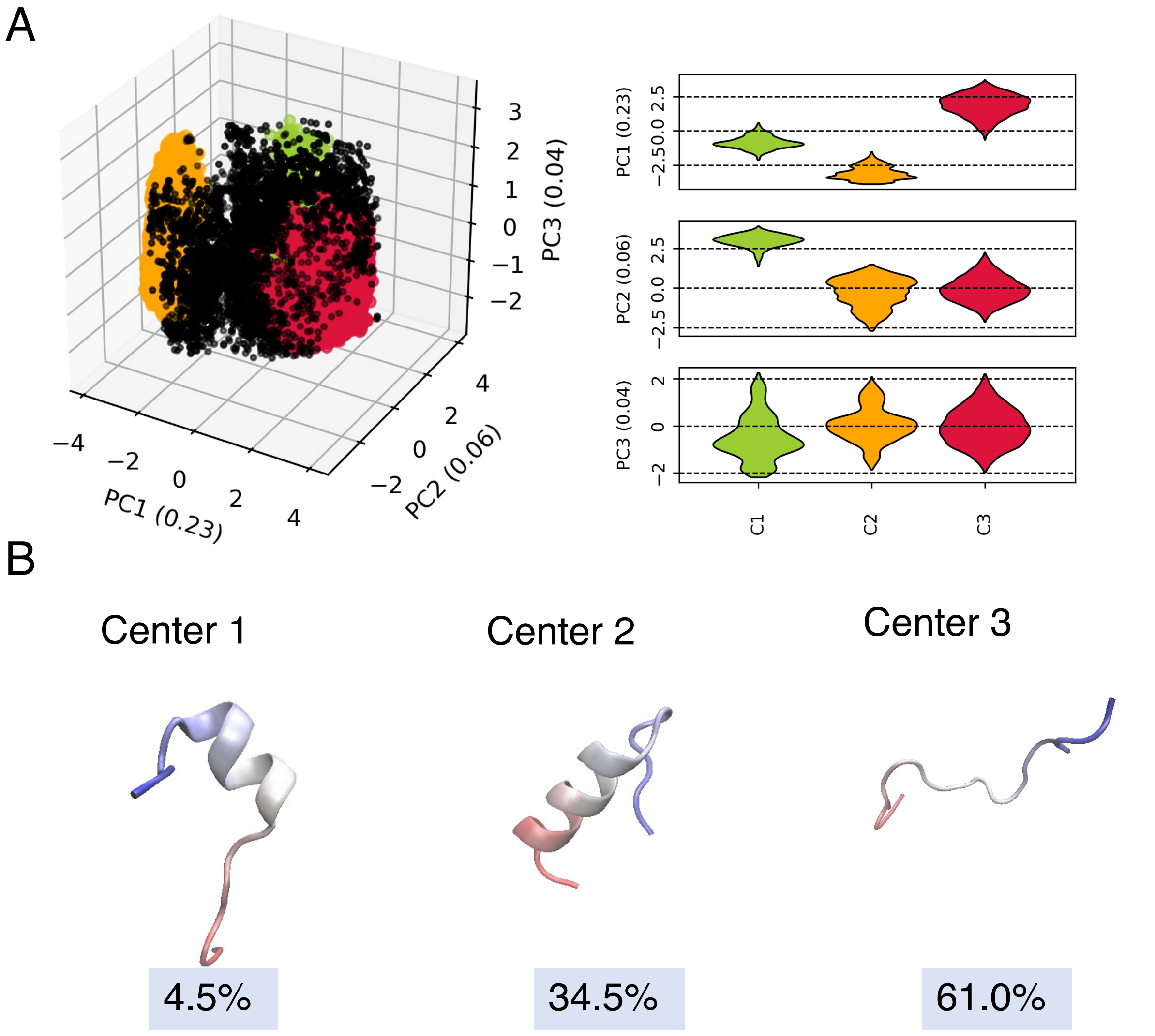


**Figure S3. Major conformation extraction from dihedral angle.** (A) Clustering result for Nt19 WT based on the first 3 PC. (B) Conformation corresponds to the center of each cluster and the abundance of each cluster. The molecules are colored red to blue from N-term to C-term.

| | C1 | C2 | C3 | C4 | C5 |
|---|---|---|---|---|---|
| WT | 4.5% | 34.5% | 61.0% | | |
| | 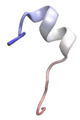 | 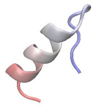 | 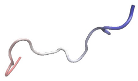 | | |
| AcK6 | 17.6% | 21.9% | 60.5% | | |
| | 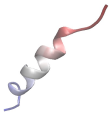 | 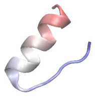 | 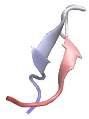 | 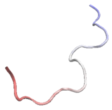 | |
| AcK9 | 7.3% | 6.2% | 20.3% | 66.2% | |
| | 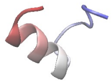 | 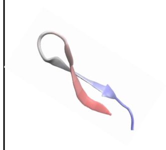 | 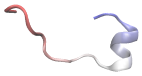 | 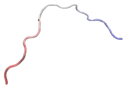 | |
| pT3 | 3.9% | 6.8% | 19.4% | 46.1% | 23.8% |
| | 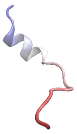 | 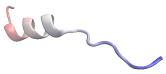 | 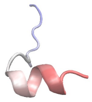 | 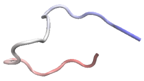 | 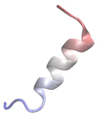 |

**Figure S4. Major conformations and corresponding abundances for WT, AcK6, AcK9, and pT3.** The major conformations for each unmodified and modified Nt19 were obtained by clustering based on dPCA of the in-solution simulations. The molecules are colored red to blue from N-term to C-term.

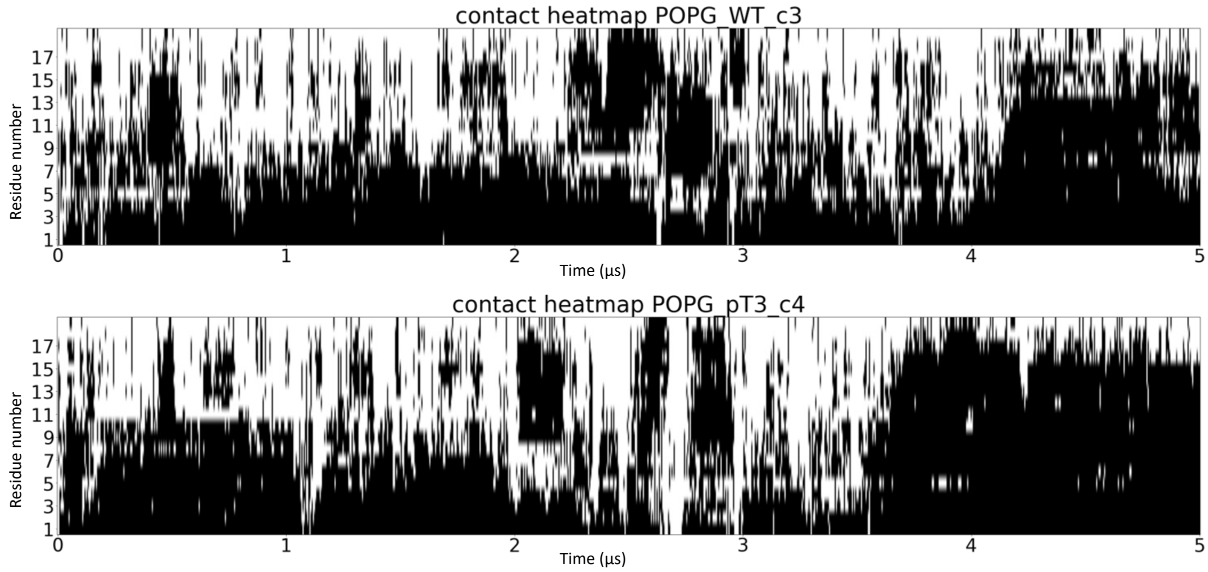


**Figure S5. The disordered conformation of pT3 had a similar affinity to the membrane compared to the unmodified Nt19.** The contact heatmap for the disordered conformation of pT3 and unmodified Nt19 showed that the disordered pT3 conformation had a similar interaction with the membrane.

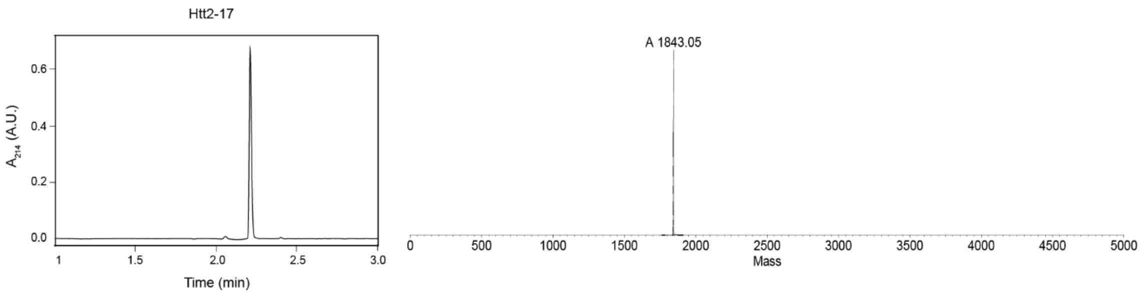
Htt2-17
A214 (A.U.)
Time (min)
A 1843.05
Mass

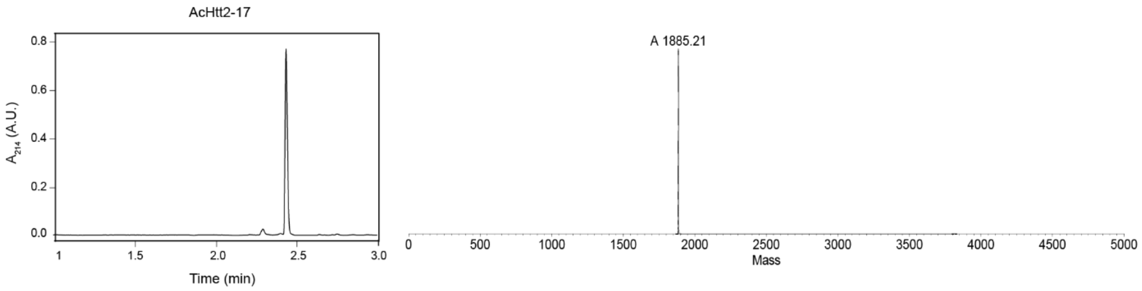
AcHtt2-17
A214 (A.U.)
Time (min)
A 1885.21
Mass

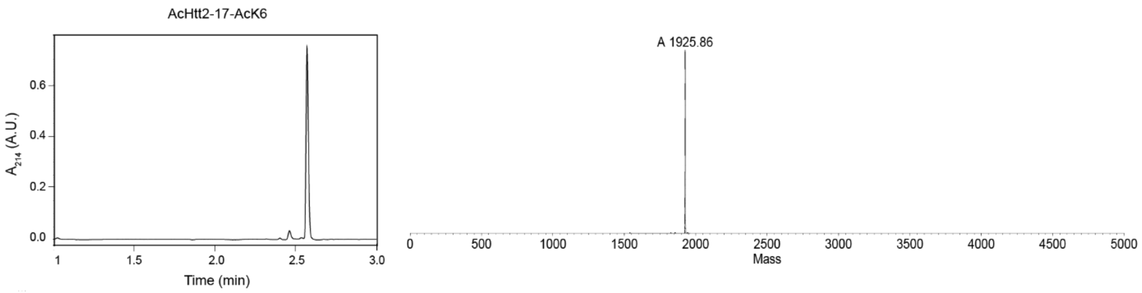
AcHtt2-17-AcK6
A214 (A.U.)
Time (min)
A 1925.86
Mass

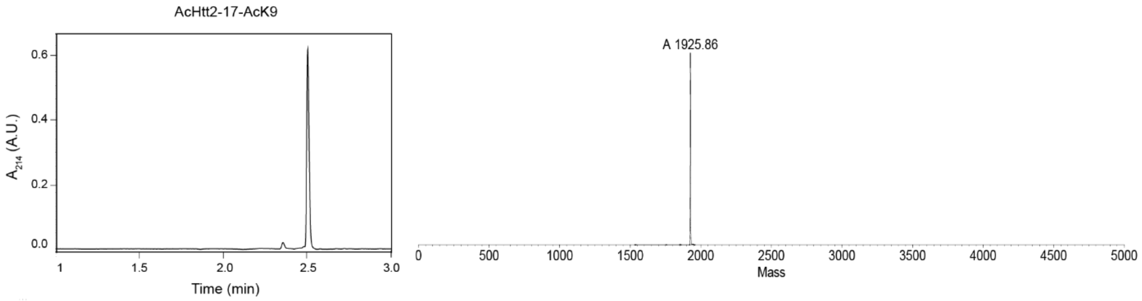
AcHtt2-17-AcK9
A214 (A.U.)
Time (min)
A 1925.86
Mass

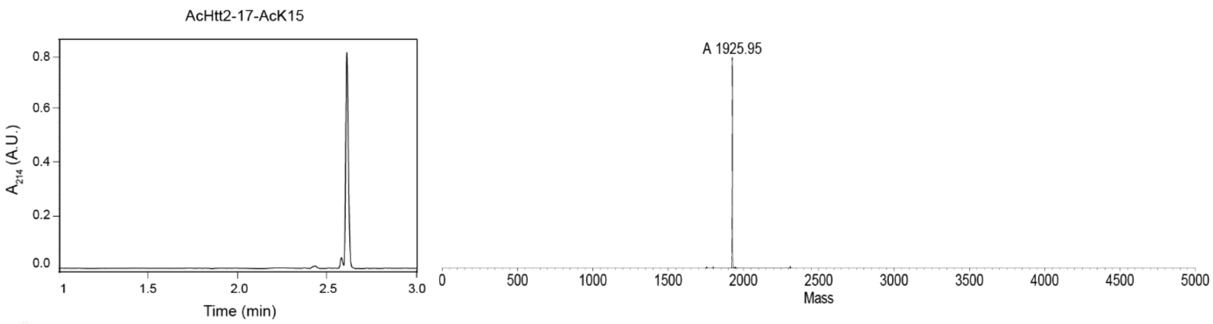
AcHtt2-17-AcK15
A214 (A.U.)
Time (min)
A 1925.95
Mass

Htt1-19

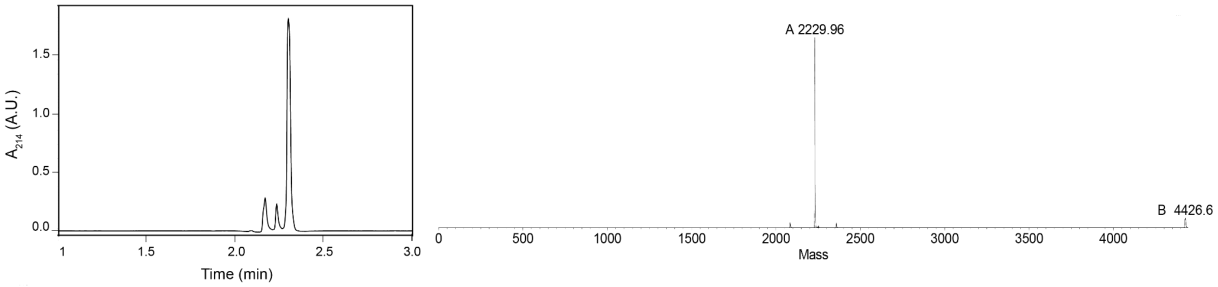


Htt1-19 pS13

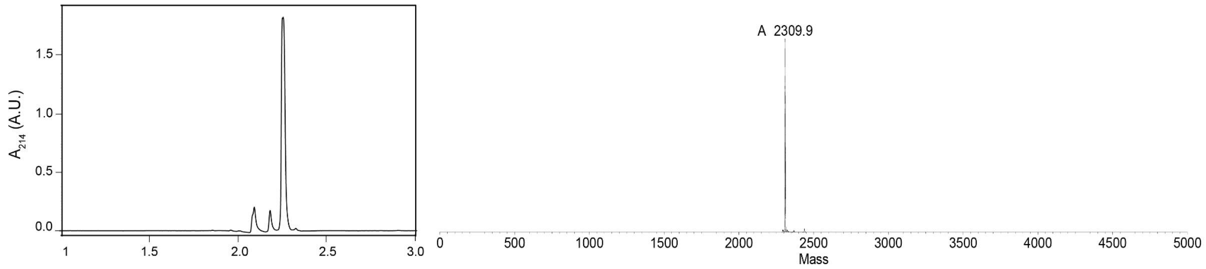


Htt1-19 pS16

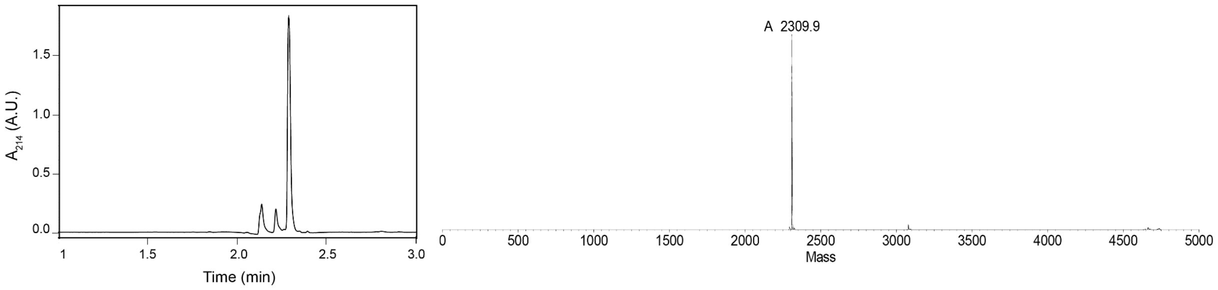


Htt1-19 pT3 pS13

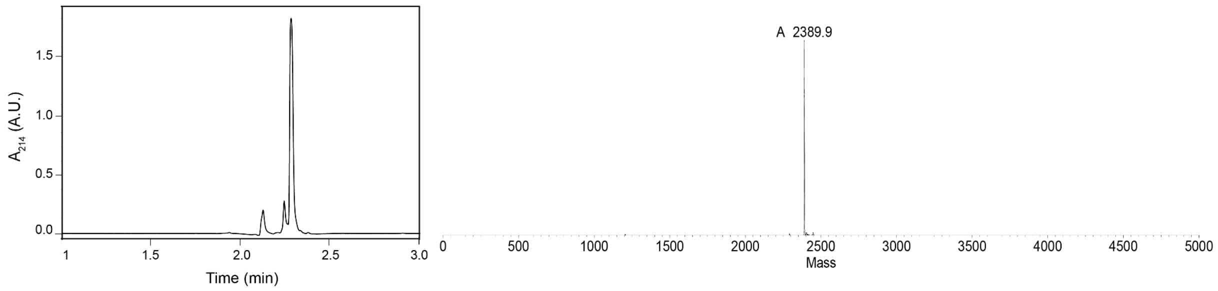

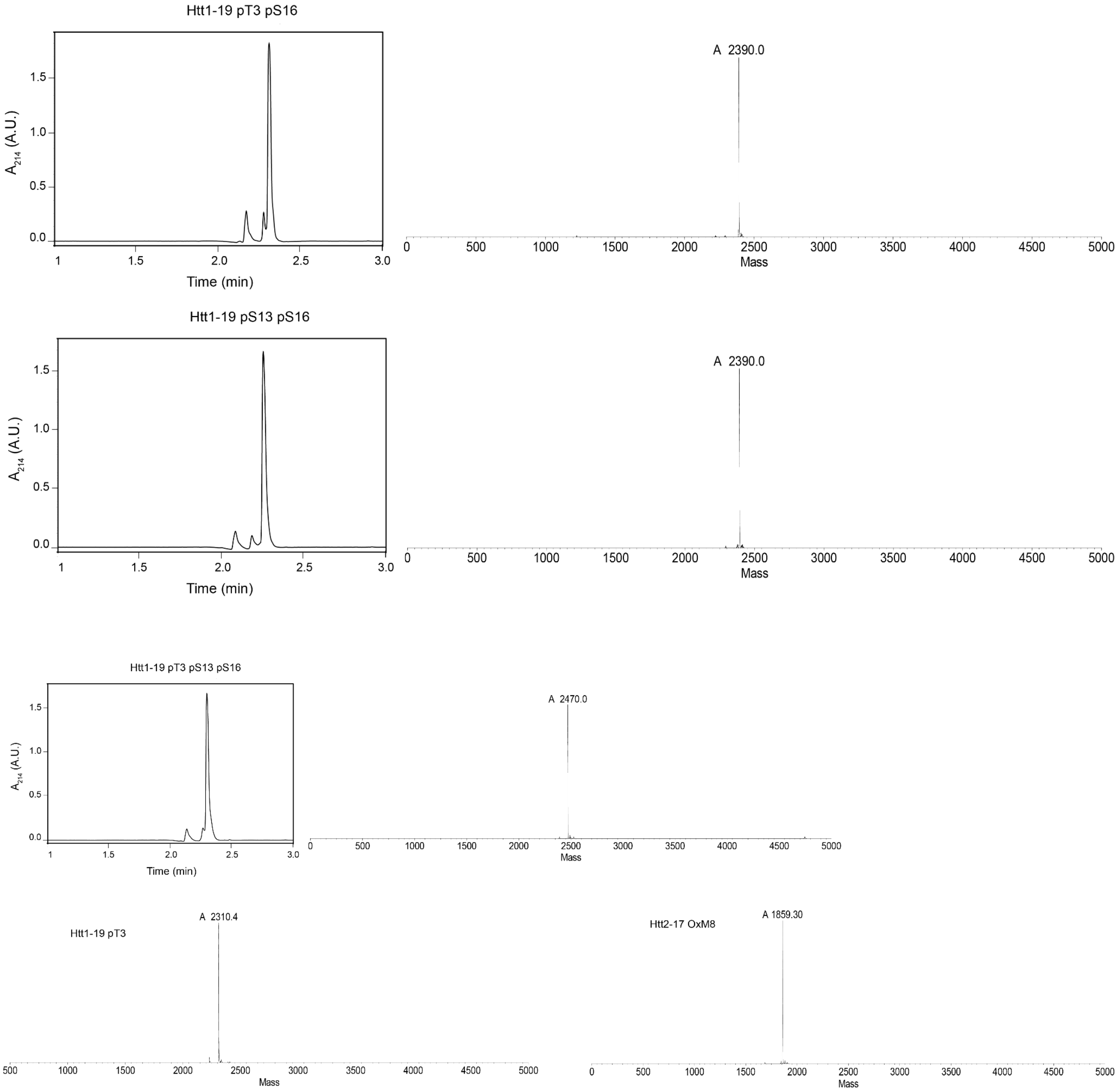


**Figure S6. Purity analysis of all peptides from this study assessed by RP-UPLC and LC-MS.**

**A**

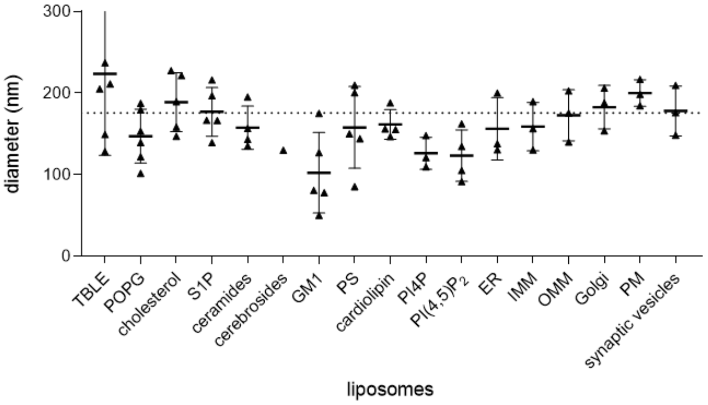


**B**

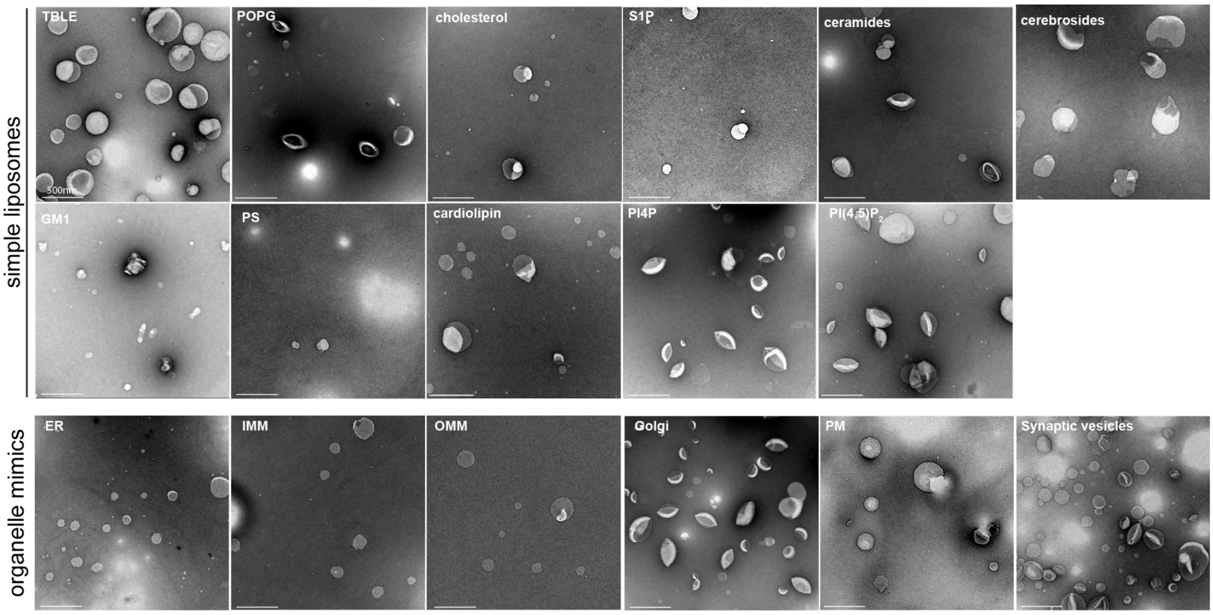


**Figure S7. Size distribution of lipid vesicles assessed by A) DLS and B) TEM.**
